# Small-molecule-mediated OGG1 inhibition attenuates pulmonary inflammation and lung fibrosis

**DOI:** 10.1101/2021.02.27.433075

**Authors:** L. Tanner, A.B. Single, R.K.V Bonghir, R. Oomen, O. Wallner, T. Helleday, C. Kalderen, A. Egesten

## Abstract

Interstitial lung diseases such as idiopathic pulmonary fibrosis (IPF) are caused by persistent micro-injuries to alveolar epithelial tissues together with aberrant repair processes. Despite substantial advancement in our understanding of IPF progression, numerous questions remain concerning disease pathology. IPF is currently treated with pirfenidone and nintedanib, compounds which slow the rate of disease progression but fail to treat underlying causes of disease. The DNA repair enzyme 8-oxoguanine DNA glycosylase-1 (OGG1) is upregulated following TGF-β exposure in several fibrosis-associated cell types. Currently, no pharmaceutical solutions targeting OGG1 have been utilized in the treatment of IPF. In this study, a novel small molecule OGG1 inhibitor, TH5487, decreased myofibroblast transition and associated pro-fibrotic markers in fibroblast cells. In addition, TH5487 decreased pro-inflammatory cytokine production, inflammatory cell infiltration, and lung remodeling in a murine model of bleomycin-induced pulmonary fibrosis. Taken together, these data strongly suggest that TH5487 is a potent, specific, and clinically-relevant treatment for IPF.

## Introduction

Idiopathic pulmonary fibrosis (IPF) is a disorder characterized by progressive lung scarring with a median survival time of 3 years post-diagnosis (1–3). The disease is associated with increasing cough and dyspnea, affecting approximately 3 million people worldwide, with incidence strongly correlated with increasing age (4). IPF is defined on the histological basis of usual interstitial pneumonia (UIP), with UIP usually presenting as ‘honeycombing’ and peripheral alveolar septal thickening (5). Fibroblast and myofibroblast overactivation/overstimulation results in extracellular matrix (ECM) deposition in alveolar walls, reducing alveolar spaces (6). Current IPF therapies focus on inhibiting collagen deposition by blocking myofibroblast activation. These approaches have shown limited success in achieving overall IPF resolution, highlighting the need for novel therapeutic strategies (4, 7).

Injured IPF tissues produce reactive oxygen species (ROS), resulting in DNA damage and the upregulation of fibrotic-related pathways, ultimately leading to lung architecture collapse (8, 9). The DNA base guanine is particularly prone to oxidation, forming 7,8-dihydro-8-oxoguanine (8-oxoG). 8-oxoG bases are recognized by the enzyme 8-oxoG DNA glycosylase 1 (OGG1), whereby OGG1 binding triggers DNA base excision processes. It has been shown that Ogg1-deficient (*Ogg1 ^-/-^)* mice are resistant to DNA damage inducing agents (10). Furthermore, the increased expression of OGG1 is an important marker of inflammation and resultant fibrotic processes (11, 12).

Following lung injury, fibroblasts transition to myofibroblasts through stimulatory factors such as TGF-β1, which further induce the production of fibrotic markers including alpha smooth muscle actin (α-SMA), collagen, and fibronectin (13, 14). The fibroblast to myofibroblast transition and migratory activities are well established hallmarks of IPF (15, 16).

Interestingly, siRNA-mediated *Ogg1* knockdown in murine embryonic fibroblast cells revealed decreased levels of tissue-associated α-SMA (17). OGG1’s implication in fibrogenesis, combined with its role in inflammation, highlights this enzyme as a therapeutic target for IPF treatment (18). Thus, we hypothesize that efficacy of a previously identified, potent, and selective OGG1 inhibitor (TH5487) may show promise in a murine model of IPF. In this study, we used a 21-day intratracheally bleomycin-challenged murine model to confirm TH5487 efficacy *in vivo*. This model reproduces several phenotypic features of human IPF, including peripheral alveolar septal thickening and acute cytokine production resolving in fibrosis (19, 20).

## Methods

### Cell culture

Murine C57BL/6 embryonic lung fibroblasts (MEF’s; ATCC, Manassa, VA) and HFL-1 cells were cultured in complete growth medium supplemented with L-glutamine, 100 U/mL penicillin, 100 µg/mL streptomycin, and 10% FBS. A549 cells were cultured in RPMI 1640 with the addition of 10% FCS, and 100U/mL penicillin, 100 µg/mL streptomycin. Finally, BEAS 2B cells were cultured in RPMI-glutamax and 10% FBS, with the addition of 100 U/mL penicillin and 100 µg/mL streptomycin.

### Wound healing assay

Wound healing assays were conducted using A549, murine lung fibroblast (MEF), BEAS 2B, and HFL-1 cells grown to confluence in 24-well plates. Wounds were made in the confluent cell layer using sterile 200 uL pipette tips, followed by washing with PBS and incubation with complete culture medium with or without TGF-β1 (10 ng/mL) at 37 °C in a 5 % CO_2_ incubator for 48 h post-scratching. Cell images were monitored every 24 h using an Olympus CKX41 microscope with Olympus SC30 camera and cellSens Entry software (Olympus, Tokyo, Japan). Images were analyzed using the wound healing tool in ImageJ. Data are presented as percentages of the initial wound area.

### Immunostaining of HFL-1 cells

HFL-1 cells were seeded (1 × 10^4^ cells/mL) into 24 well plates containing rounded glass cover slips. TGF-β1 (10 ng/mL) was added to each well, followed by the addition of medium containing TH 5487, dexamethasone, medium only, or vehicle only. Following 96 h of treatment, cells were washed and fixed with ice cold methanol, containing 0.5% Triton-X100. Slides were blocked using Dako Protein Block (Agilent, CA, USA) for 1 h at room temperature and then stained with primary antibodies overnight, rabbit anti-COL I, rabbit anti-αSMA, rabbit anti-fibronectin (Abcam, CAM, UK), and rabbit anti-OGG1 (Invitrogen, Carlsbad, CA) antibodies were used. AlexaFluor 488-conjugated goat anti-rabbit antibody (Invitrogen, CA, USA) was used as secondary antibody. Glass cover slips were removed and mounted onto glass slides, with nuclei counterstained using DAPI-containing fluoroshield (Abcam). Images were visualized using a Nikon Confocal Microscope.

### Fibroblast transwell experiment

Fibroblast chemotaxis was measured using 24-well Nunc (8 µm pore size) transwell inserts (ThermoFisher, MA, USA). HFL-1 cells were seeded (5 × 10^5^ cells/ml) into the upper chamber in FBS-free medium, whilst the lower chamber contained complete medium with additional 10 % FBS as a chemoattractant. Medium containing TGF-β1 (10 ng/mL) was added to each well and allowed to equilibrate for 24 h. Cells were washed with PBS, followed by the addition of medium containing TH5487, dexamethasone, medium only, or vehicle only (DMSO 2% and PBS pH 7.4), with the replacement of each every 24 h. Following 96 h, medium was removed and cells in the lower chamber were stained (crystal violet) and imaged using a Nikon microscope with a x10 objective.

### mRNA analysis

HFL-1 cells were starved in culture medium with 0.5% BSA for 24 h. Cells were pretreated with TH5487, 5 μM, 10 μM or DMSO for one hour followed by TGF-β stimulation (10 ng/ml, Peprotech) for 48 hours. mRNA was extracted based on the RNA extraction kit protocol Direct-zol RNA MiniPrep (Zymo research).

cDNA synthesis was performed using Quantitect Reverse Transcription kit (Qiagen) and quantified with the iTaq™ Universal SYBR® Green Supermix PCR Kit in Rotor Gene Q (Qiagen). Using the following pcr primers, Human Col1A1F, 5’ GATTCCCTGGACCTAAAGGTGC3’, Human Col1A1R, 5’ AGCCTCTCCATCTTTGCCAGCA3’, Human α-SMAF, 5’ CTATGCCTCTGGACGCACAACT3’, Human α-SMAR 5’ CAGATCCAGACGCATGATGGCA3’, 18SrRNAF: 5’ AGTCCCTGCCCTTTGTACACA 3’, 18SrRNAR: 5’ GATCCGAGGGCCTCACTAAAC 3’

The obtained Ct values of the samples were normalized to the Ct values of 18SRNA reference gene, 2-ΔΔCt values were calculated using Microsoft Excel.

### Ethical approval

All animal experiments were approved by the Malmö-Lund Animal Care Ethics Committee (M17009-18).

### Animals

10-12-week-old male C57Bl/6 mice (Janvier, Le Genest-Saint-Isle, France) were housed at least 2 weeks in the animal facility at the Biomedical Service Division at Lund University before initiating experiments and were provided with food and water *ad libitum* throughout the study. Mice were randomly allocated into five groups: intratracheally (i.t.)-administered bleomycin (2.5 U/kg) + vehicle intraperitoneal (i.p.), bleomycin (i.t.) + TH5487 (40 mg/kg; i.p.), bleomycin (i.t.) + dexamethasone (10 mg/kg; i.p.), saline (i.t.) + vehicle (i.p.), and saline (i.t.) + TH5487 (40 mg/kg; i.p.).

### Blood collection

Blood was collected in 0.5 M EDTA tubes by cardiac puncture and centrifuged at 1,000 g for 10 min. Plasma supernatant was used for the analysis of inflammatory mediators using a bioplex assay.

### Collection of lung tissue

Right lungs were collected in Eppendorf tubes on dry ice and stored at -80 °C. The snap-frozen lungs were thawed and homogenized in tissue protein extraction reagent (T-PER) solution (ThermoFisher) containing protease inhibitor (Pefabloc SC; Sigma-Aldrich) at a final concentration of 1 mM. Lung homogenates were centrifuged at 9,000 g for 10 min at 4°C, and the supernatants were collected for bioplex analysis. Left lungs were collected in Histofix (Histolab, Göteborg, Sweden) and submerged in 4 % buffered paraformaldehyde solution.

### Bronchoalveolar lavage fluid (BALF) collection

BAL was performed with a total volume of 1 ml PBS containing 100 μM EDTA. BALF was collected in Eppendorf tubes on ice, with aliquots made for flow cytometry, cytospin differential counts, and an aliquot transferred to -80°C for bioplex cytokine analysis. Cytospin preparations of cells were stained with modified Wright-Giemsa stain (Sigma-Aldrich, St. Louis, MO).

### Bioplex cytokine analysis

For the detection of multiple cytokines in BALF, plasma, and lung homogenate, the Bio-Plex Pro mouse cytokine assay (23-Plex Group I; Bio-Rad) was used on a Luminex-xMAP/Bio-Plex 200 System with Bio-Plex Manager 6.2 software (Bio-Rad, Richmond, CA). A cytometric magnetic bead-based assay was used to measure cytokine levels, according to the manufacturer’s instructions. The detection limits were as follows: Eotaxin (4524.58-1.23 pg/mL), GCSF (99318.6-7.3 pg/mL), GMCSF (6310.48-3.91 pg/mL), IFN-γ (16114.01-0.87 pg/mL), IL-1α (10055.54-0.54 pg/mL), IL-1β (31512.04-1.75 pg/mL), IL-2 (19175.48-1.24 pg/mL), IL-3 (7514.5-0.44 pg/mL), IL-4 (5923.58-0.34 pg/mL), IL-5 (12619.59-0.78 pg/mL), IL-6 (9409.63-0.68 pg/mL), IL-9 (64684.09-2.41 pg/mL), IL-10 (77390.75-4.18 pg/mL), IL-12p40 (144560.15-18.62 pg/mL), IL-12p70 (78647.56-4.81 pg/mL), IL-13 (197828.67-11.16 pg/mL), IL-17 (8727.85-0.51 pg/mL), KC (23001.9-1.4 pg/mL), MCP-1 (393545.52-10.01 pg/mL), MIP-1α (14566.62-0.63 pg/mL), MIP-1β (7023.87-0.34 pg/mL), RANTES (19490.48-4.61 pg/mL), and TNF-α (74368.54-51.69 pg/mL). Cytokine measurements for lung homogenate samples were corrected for total protein concentration using a Pierce™ BCA Protein Assay Kit (ThermoFisher).

### TGF-β1 ELISA

The Quantikine ELISA kit targeting TGF-β1 (R&D systems, UK) was used to assess TGF-β1 levels in the BALF, plasma, and lung homogenate of murine samples according to the manufacturer’s instructions. Optical density was measured at 450 nm using a VICTOR 1420 Multilabel plate reader (Perkin Elmer, MA, USA).

### Flow cytometry

Flow cytometry was carried out using a BD Accuri C6 Plus (BD Biosciences). The washed cells were incubated with Fixable Viability Stain 510 (FVS510; BD #564406) to differentiate live and dead cells. Cells were washed with Stain buffer 1x (BD #554656) and incubated with Lyse Fix 1x (BD #558049 (5x)). Fixed cells were washed with stain buffer and aliquoted into two samples incubated with either anti-CD11b (BD553312), anti-CD11c (BD558079), anti-Ly6G (BD551461), or anti-CD11c, anti-MHCII (BD558593), anti-SiglecF (BD562680) antibodies.

### H&E and picrosirius red staining of lung tissue

Mouse left lungs were fixed in Histofix (Histolab Products AB, Askim, Sweden), paraffin-embedded and sectioned at 3 µm. The tissue sections were placed on slides (Superfrost Plus; Fisher Scientific) and deparaffinized in serial baths of xylene and ethanol followed by staining using Mayer hematoxylin and 0.2% eosin (Histolab Products AB, Askim, Sweden) or picrosirius red staining kit (Abcam, Cambridge, UK). The stained slides were imaged using an Aperio CS2 image capture device.

### SEM analysis of lung histology

Lung tissue sections were fixed as reported above. After fixation, samples were washed and dehydrated in alcohol at increasing concentrations, critical point dried, mounted on aluminium holders, and covered with 20 nm of gold. Samples were examined in a Philips XL30 FEG scanning electron microscope (Eindhoven, The Netherlands) operated at an acceleration voltage of 5 kV.

### Immunostaining of murine lung sections

Lung tissue sections were fixed as reported above. Lung samples underwent antigen retrieval (pH9 buffer) using a Dako PT Link pre-treatment module (Agilent, CA, USA). Samples were washed and blocked for 10 min (Dako protein block, Agilent, CA, USA) before being treated with primary antibodies overnight. Mouse anti-COL1A1, rabbit anti-fibronectin, rabbit anti-MPO (Abcam, CAM, UK), and rabbit anti-OGG1 (Invitrogen, Carlsbad, CA) antibodies were used. Alexa Fluor 488-conjugated goat/anti-mouse and Alexa Fluor 647 goat/anti-rabbit (Invitrogen, CA, USA) were used as secondary antibodies. Glass cover slips were placed onto slides and mounted with DAPI-containing fluoroshield (Abcam). Images were visualized using a Nikon Confocal Microscope.

## Results

### TH5487 reduced migratory capacity of lung-resident cells

The migration of fibroblasts into the lung injury site is a key step in the pathogenesis of pulmonary fibrosis. The human HFL-1 and MEF cell lines were used for this analysis, alongside the alveolar epithelium-derived cell lines A549 and BEAS2B. Cells were grown to form a confluent monolayer, scratched to form a wound, then stimulated with 10 ng/mL TGF-B1 to induce cell migration and wound closure. The cells were additionally treated with TH5487 (10 uM), vehicle, or dexamethasone (10 uM) as a comparator drug. This assay showed the wound area percentage in all cell types was significantly reduced following TGF-β1 addition, indicating the expected pro-migratory effects of TGF-B1 stimulation (Figure 1A). TGF-B1-induced wound closure was inhibited by TH5487 treatment across all tested cell lines (Figure 1A). Interestingly, dexamethasone only inhibited cell migration in BEAS 2B and HFL-1 cells and showed minimal effects in A549 and mouse lung fibroblast cells. In summary, these results indicate TH5487 is capable of inhibiting TGF-B1-mediated migratory effects of lung-derived cells, including both human- and mouse-derived fibroblasts.

**Figure 1:**
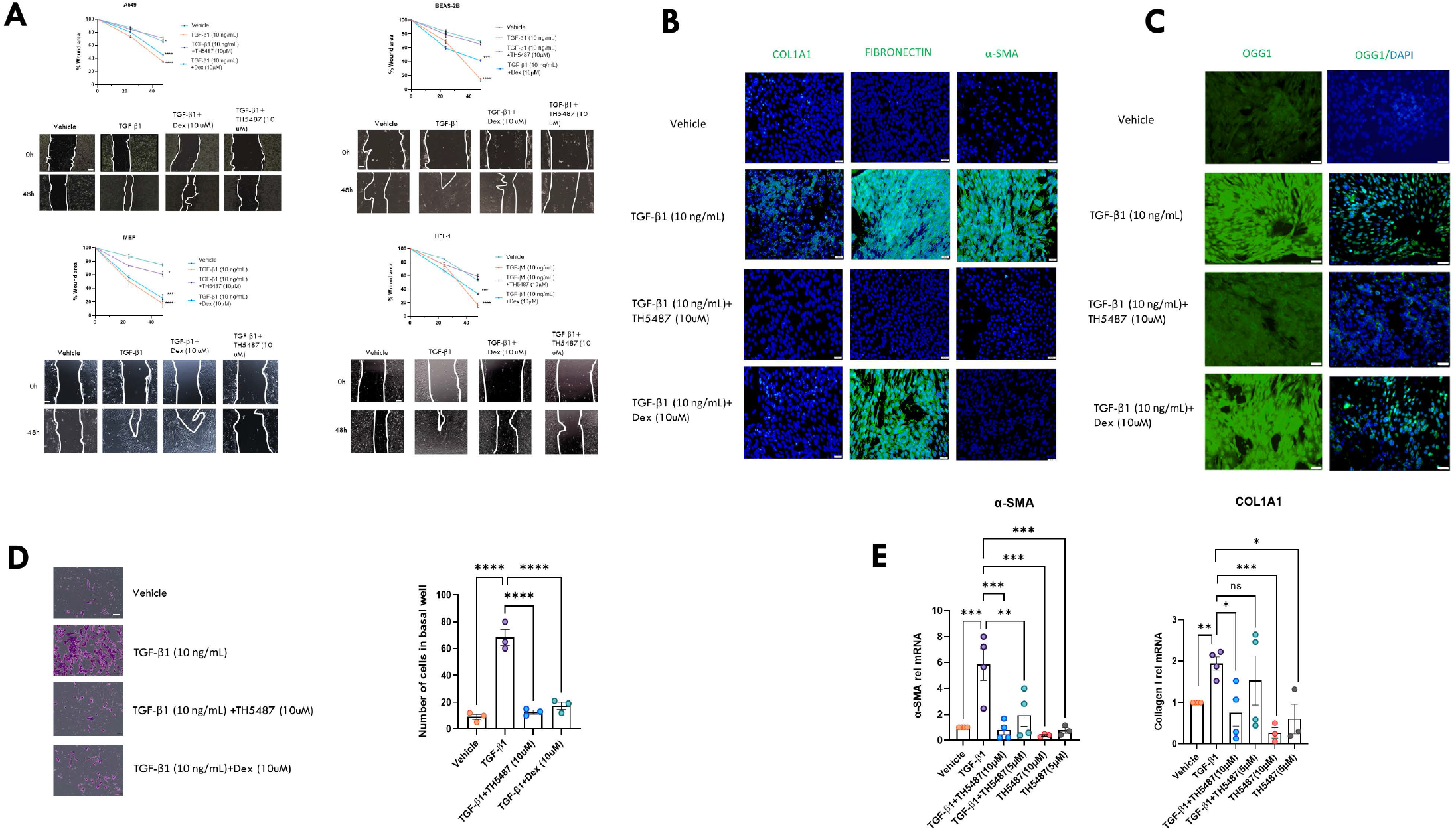
*In vitro* wound healing assay (A), immunofluorescence and myofibroblast transition assay using human lung fibroblast (HFL-1) cells (B-D), and qPCR of HFL-1 cells following TGF-β1 stimulation. (A) Migration of A549, mouse lung fibroblast (MEF), BEAS2B, and human lung fibroblast cells (HFL-1) was measured at 0, 24, and 48 h post-wound induction. Cells were treated with 10 ng/mL of TGF-β1 or an equivalent concentration of DMSO in control samples. Wound area percentage was compared to the control samples using a one-way ANOVA followed by a Dunnett’s post-hoc test: *P<0.05; ***P<0.0005 ****P<0.0001. Data is representative of 3 independent experiments containing 4 biological replicates (scale bar=100 μm). Immunostaining of myofibroblast cells (B and C) following 96h of TGF-β1 treatment, with TH5487 treatment displaying visually reduced levels of collagen (COL1A1), fibronectin, and alpha smooth muscle actin (α-SMA) with similarly reduced levels of OGG1 compared to no treatment/ TGF-β1 control (green fluorescence; scale bar=50 μm). (D) Myofibroblast transition of human lung fibroblast cells was measured over 96h. HFL-1 cells in the basal well were stained and counted, with significantly more myofibroblast cells appearing in the untreated control well. Drug treated experiment samples were compared to untreated control samples using a one-way ANOVA followed by a Dunnett’s post-hoc test: ****P<0.0001. Data is representative of 3 independent experiments containing 4 biological replicates (scale bar=100 μm). (E) The effects of TH5487 on transcription of α-SMA and Col1A1 in TGF-β1 stimulated HFL-1 cells were investigated by qPCR. TH5487 (10 μM), significantly reduced transcription of ColA1 and α-SMA, with data representative of 3 independent experiments.

### TH5487 reduced HFL-1 myofibroblast transition, whilst targeting OGG1 production

TGF-β1 driven fibrosis results in the transition of fibroblasts to myofibroblasts resulting in the production of fibrotic proteins. TH5487 treatment additionally reduced the production of collagen, fibronectin, and α-smooth muscle actin, as measured by immunofluorescence and qPCR (Figure 1B and E). Compelling evidence of TH5487 involvement in OGG1 reduction is presented in Figure 1C, with significant decrease in OGG1 fluorescence compared to TGF-β1. To investigate myofibroblast transition, TGF-B1-stimulated HFL-1 cells were analyzed for TH5487-mediated inhibition to myofibroblasts in a transwell assay (Figure 1D). Interestingly, TH5487 significantly reduced this myofibroblast transition (*P* <0.001) compared to TGF-β1-stimulated HFL-1 cells.

### Intraperitoneal TH5487 maintained murine weight following bleomycin administration

We next investigated TH5487-mediated Ogg1 inhibition *in vivo* using a mouse model of pulmonary fibrosis. Mice received a one-off intratracheal administration of bleomycin (2.5 U/kg) and were dosed (IP) with TH5487 or dexamethasone 1h post-bleomycin administration, followed by additional dosing once per day, in five-day intervals, over a total period of 21 days (Figure 2A). Body weights were recorded as a proxy for murine health status (Figure 2B). Mice in the bleomycin/vehicle group lost 26.5 ± 9.1 % total bodyweight, whilst mice in remaining groups either maintained their bodyweight or gained significant weight compared to the bleomycin/vehicle group over the experiment timeline.

**Figure 2:**
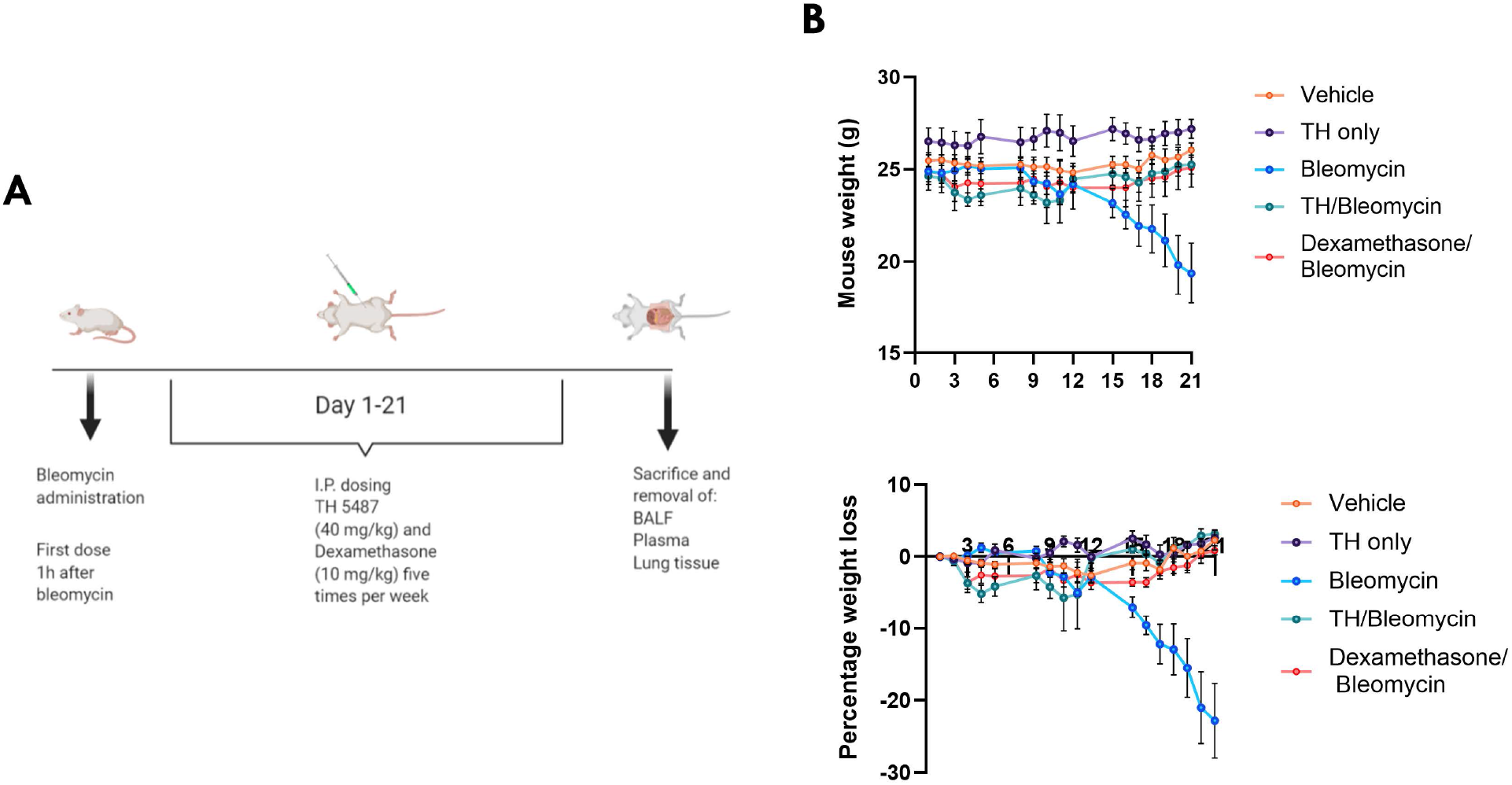
TH5487 murine dosing strategy and weights. **(A)** Mice received intratracheally administered bleomycin (2.5 U/kg) and were subsequently dosed (IP) with TH5487 or dexamethasone 1h post-bleomycin administration. Intraperitoneal (i.p.) dosing occurred five times per week, over the course of 21 days, followed by euthanasia and removal of BALF, plasma, and lung tissues. (B) Mice receiving bleomycin showed weight loss up until day 10, where after those dosed with TH5847 picked up significant amounts of weight compared to the vehicle/bleomycin group.

### TH5487 treatment reduced cytokine levels *in vivo*

The effect of TH5487 treatment on inflammatory cytokine levels was investigated from the *in* vivo study using a 23-cytokine multiplex assay (Supp. Figure 1-3). Cytokines were assessed in lung tissue homogenate, BALF, and plasma (Figure 3 A-C). TH5487 treatment significantly reduced the levels of several bleomycin-induced inflammatory cytokines, including those in the BALF such as IL-9, eotaxin, MIP-1α (\**P*<0.05), MIP-1β and IL-5 (\*\**P*<0.01), and KC (\*\*\**P*<0.005), whilst cytokines from the plasma displaying a reduction include IL-5 and MIP-1β (\**P*< 0.05), IL-17 and RANTES (\*\**P*< 0.01), and eotaxin (\*\*\*\**P*<0.001). Further, similar reductions in the lung homogenate were seen following TH5487 administration in particular in MIP-1α, KC, and IL-9 (\**P*< 0.05) and IL-2 (\*\**P*< 0.01). Additionally, TGF-β1 levels were measured in the BALF, plasma, and lung homogenate of murine samples (Figure 3D). TH5487-treated samples displayed significantly less TGF-β1 than in vehicle/bleomycin samples as compared by one-way ANOVA (\**P<0.05; **P<0.01; ***P<0.005*). Plasma leakage into the BALF as a proxy for lung damage was assessed by BCA assay (Figure 3E). TH5487-treated samples displayed significantly lower albumin levels compared to vehicle/bleomycin samples as assessed by one-way ANOVA (*\*\*P<0.01; ***P<0.005*).

**Figure 3:**
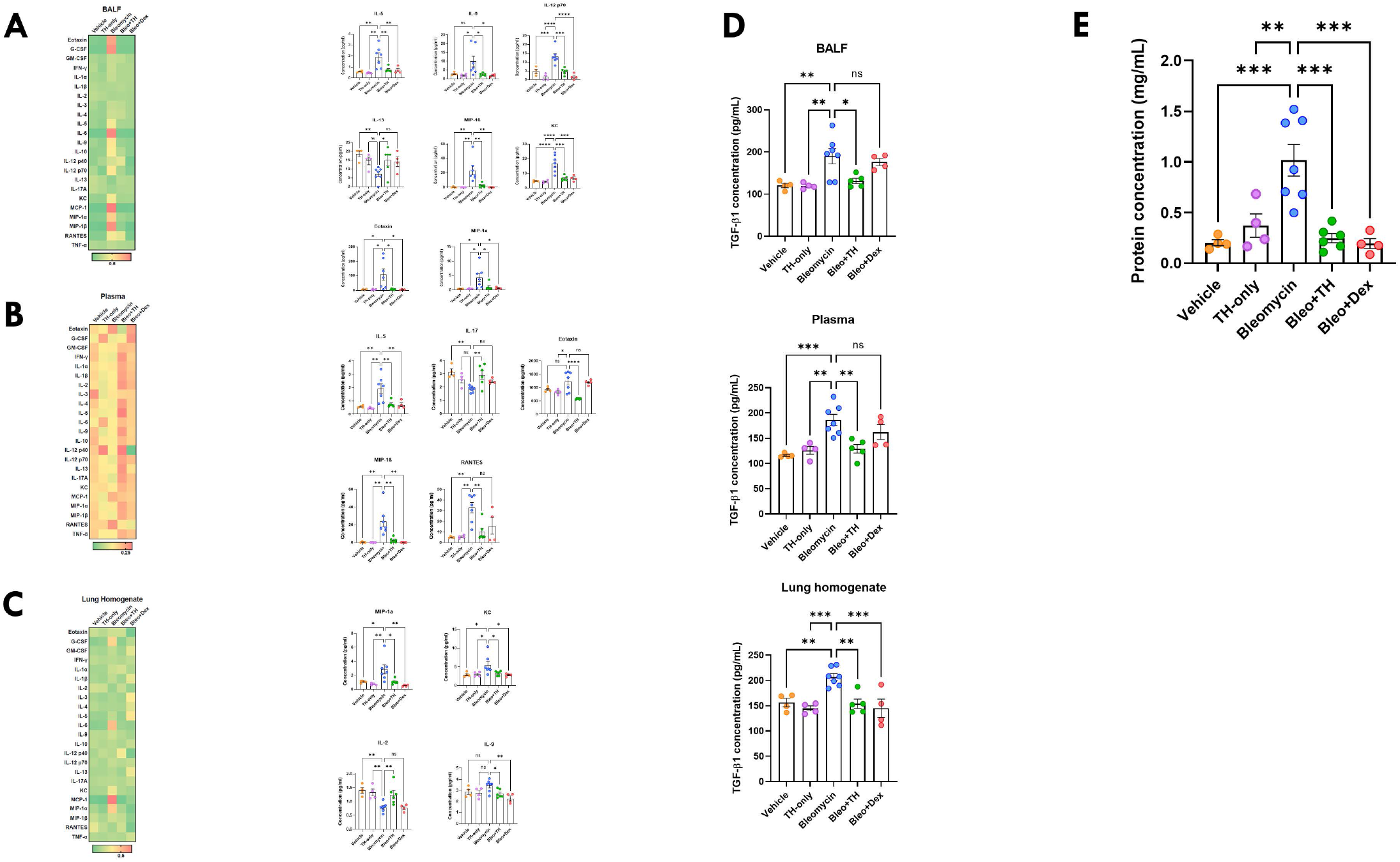
Significantly decreased murine cytokines following TH5487 (IP) administration, with accompanying markers of lung damage. Heatmaps (A-C) showing the differences in cytokine levels in murine BALF, plasma, and lung homogenate (Red-high value; Green-low value) with corresponding significantly decreased or increased cytokine levels alongside. Cytokine values were compared to the vehicle/bleomycin group using a one-way ANOVA (\**P<0.05; **P<0.01; ***P<0.005; *****P<0.001*). TGF-β1 ELISA (D) conducted on murine BALF, plasma, and lung homogenate revealed significantly decreased TGF-β1 levels in all three murine sample types with values compared to the vehicle/bleomycin group using a one-way ANOVA (\**P<0.05; **P<0.01; ***P<0.005*). Murine albumin content (E) was measured in BALF samples as a marker for plasma leakage, with TH5487 treatment significantly decreasing albumin content in the BALF compared to the vehicle/bleomycin group.

### TH5487 treatment reduced immune cell infiltration into the airways *in vivo*

We next investigated the effects of TH5487 treatment on immune cell infiltration into the airways by performing flow cytometry on BALF fluid obtained from the *in vivo* studies. Decreases in neutrophil count and inflammatory macrophages were seen following TH5487 administration compared to vehicle/bleomycin samples (Figure 4A and B). Giemsa-Wright-stained BALF samples showed bleomycin-treated macrophages were enlarged, displaying an inflammatory phenotype not seen in mice treated with TH5487 (Figure 4C).

**Figure 4:**
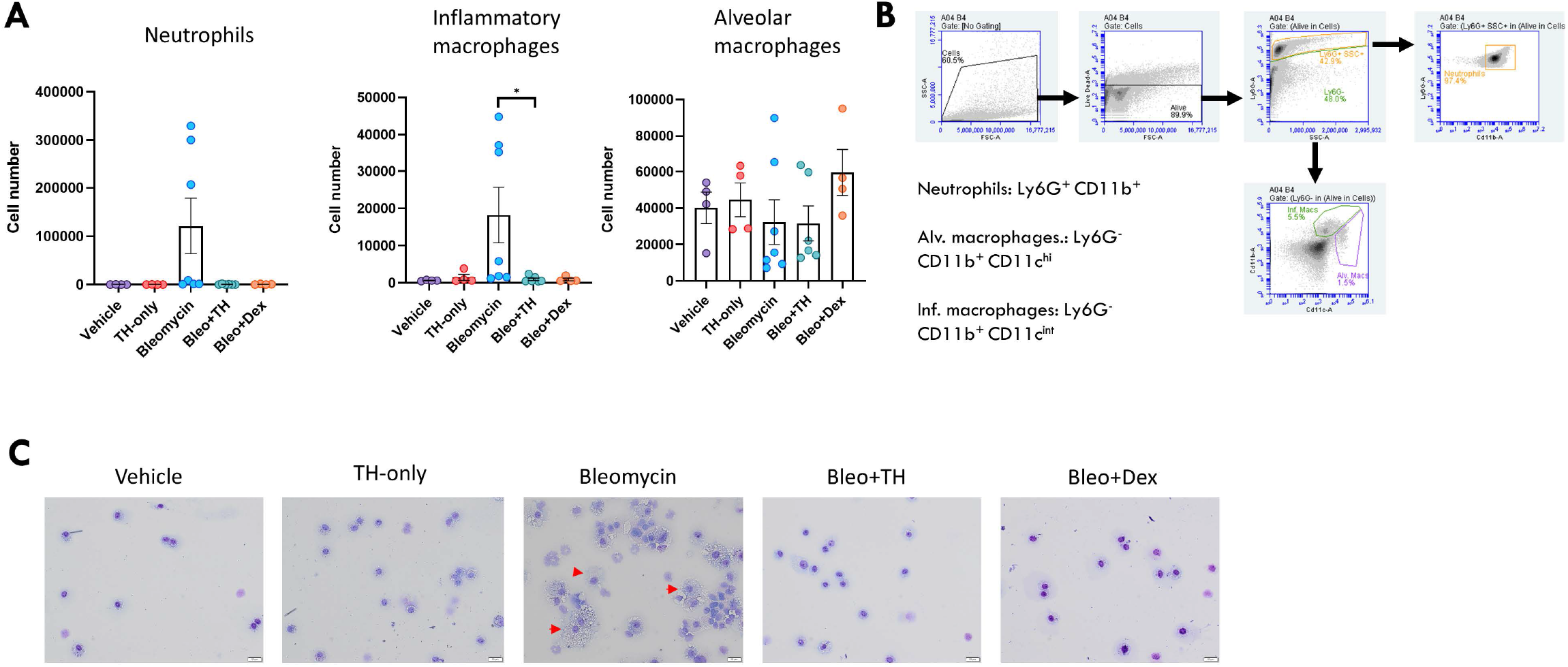
Inflammatory cell influx measured in murine BALF. Murine BALF was assessed for neutrophils, alveolar macrophages, and inflammatory macrophages (A) using flow cytometry, with the representative gating strategy depicted alongside (B). Decreased numbers of neutrophils and inflammatory macrophages were detected in response to TH5487 (IP) treatment, with no significant difference reported between the neutrophils and inflammatory macrophages of mice treated with dexamethasone. No significant differences were reported for alveolar macrophage numbers. Inflammatory cell numbers were compared to the vehicle/bleomycin group using a one-way ANOVA (\**P<0.05*). (C) Cytospin Giemsa-Wright stained slides showing vehicle/bleomycin BALF samples containing enlarged inflammatory macrophages, with TH5487 treatment reducing the presence of inflammatory macrophages, whilst the corticosteroid dexamethasone similarly reduced inflammatory macrophage influx comparable to (G) vehicle treated control samples. Scale bar=20 μm.

### TH5487 treatment decreased bleomycin-induced lung damage *in vivo*

Next, TH5487’s impact on bleomycin-induced lung damage was investigated using histological analysis of whole lung sections obtained from the *in vivo* studies (Supp. Figure 4 and 5). Bleomycin-treated control mice displayed significant lung damage when compared to saline-treated controls (Figure 5A and B). Bleomycin-treated mice that received TH5487 treatment showed reduced lung damage, with lesser degrees of alveolar structural collapse, less septal thickening of the alveoli, and less immune cell influx as indicated by the H&E stain (Figure 5A). Picrosirius red staining revealed a reduced level of collagen deposition in TH5487-treated versus control-treated mice (Figure 5B). Positive pixel analysis was used to quantify lung damage and showed significantly less H&E and picrosirius red staining in the TH 5487-administered lungs, with no significant decreases in either staining reported for dexamethasone-treated lungs (Figure 5A and B). Further, scanning electron microscopy analysis (SEM) revealed collagen deposition surrounding the alveolar walls of bleomycin control mice was reduced in response to TH5487 treatment (Figure 5C). Immunofluorescent staining of murine lung sections revealed decreased myeloperoxidase (MPO), fibronectin, COL1A1, and OGG1 staining compared to the vehicle bleomycin control group (Figure 5D). To further support the involvement of OGG1 in fibrotic related lung damage, co-staining was carried out on murine lung sections, revealing increased COL1A1/OGG1 fluorescence in similar lung areas (Figure 5E).

**Figure 5:**
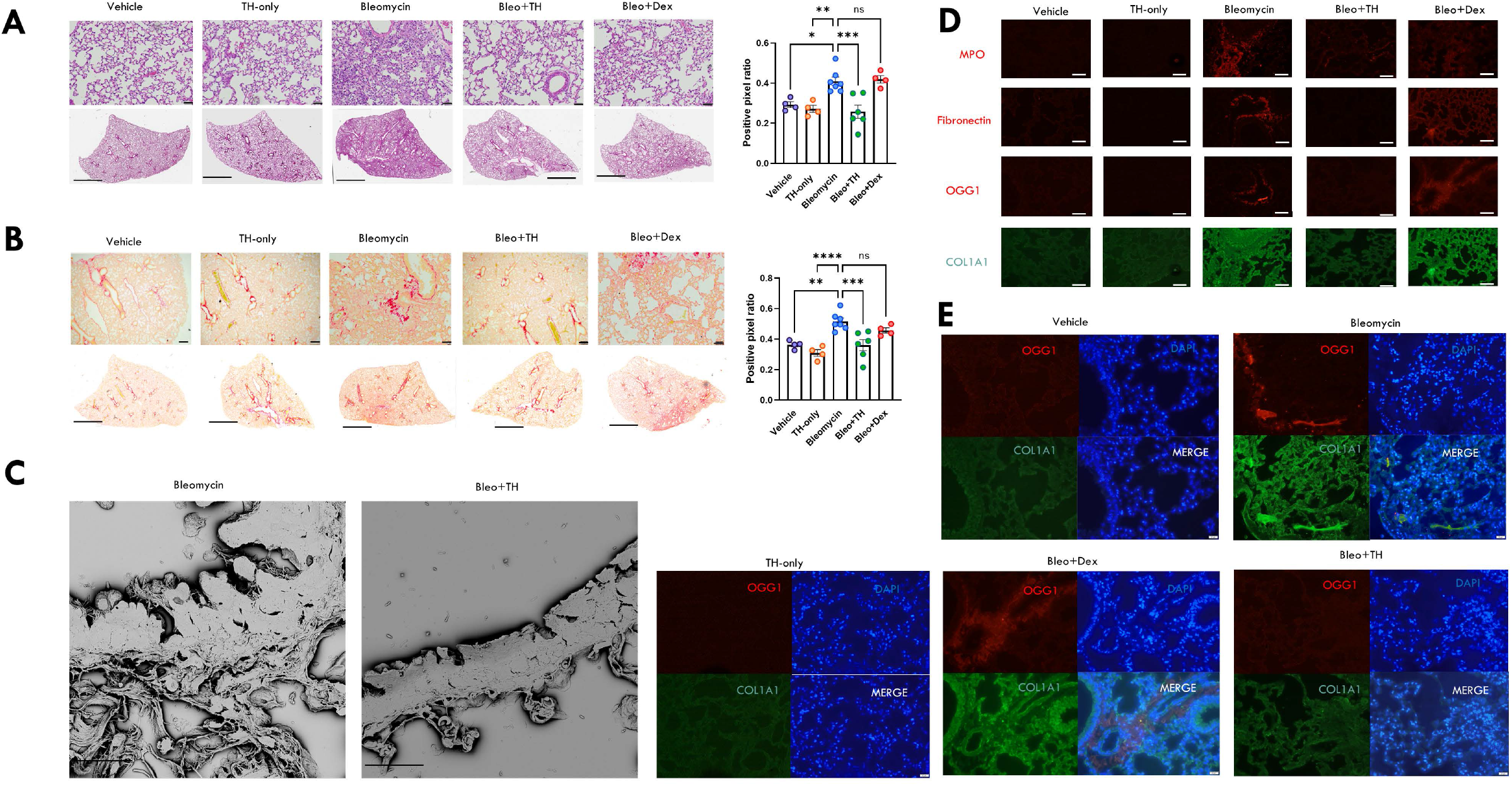
Murine lung staining, scanning electron microscopy (SEM), and immunofluorescence show reduced levels of fibrotic-related lung damage following TH5487 treatment compared to vehicle/bleomycin samples. (A) TH5487 significantly decreased lung damage (H&E) and (B) collagen deposition (picrosirius red) in both macroscopic and microscopic structures compared to vehicle/bleomycin lungs and was confirmed by positive pixel analysis of whole-lung scanned images (scale bar of microscopic image=100 μm; scale bar of whole lung scan=2 mm). Statistical analyses were conducted using a one-way ANOVA (\**P<0.05; **P<0.01; ***P<0.005*). (C) TH5487 (i.p.)/bleomycin SEM images show reduced collagen deposition in the alveolar borders compared to bleomycin-treated controls (scale bar=20 μm). Immunofluorescent staining of murine lung slices (D) revealed decreased levels of myeloperoxidase (MPO), fibronectin, OGG1, and COL1A1 following TH5487 treatment compared to both vehicle/bleomycin and dexamethasone/bleomycin groups (scale bar=50 μm). (E) Co-stained murine lung samples revealed corresponding increases in OGG1 and COL1A1 following bleomycin administration, with reduced levels of both OGG1 and COL1A1 in TH5487-treated samples (scale bar=50 μm).

## Discussion

IPF is an interstitial lung disease characterized by dysregulated inflammation, progressive lung scarring, and eventual death due to respiratory complications (1, 21). Importantly, uncontrolled lung injury is a hallmark of IPF initiation and progression, resulting in pro-inflammatory and pro-fibrotic cytokine release driving further fibrosis-related immune cell influx and ECM remodeling (16, 22, 23). Ensuing TGF-β production following lung injury results in the upregulation of tissue repair genes, including the DNA base excision repair gene OGG1 (14, 24, 25). Previously reported data indicate that OGG1 is increased in lung epithelial cells and fibroblasts following TGF-β stimulation addition (10).

Although studies showing *in vitro* and *in vivo* involvement of OGG1 in various pathologies, this is to the best of our knowledge, the first reported evidence of OGG1-targeting using a clinically relevant pharmacological agent. The approach reported in this study utilizes a potent and selective small molecule employing a novel and distinct mechanism of action, preventing OGG1 from binding damaged DNA, initiating transcription factors, and upregulating pro-inflammatory and pro-fibrotic pathways (18).

Abnormal wound healing of the alveolar epithelium in response to micro-injuries plays a crucial role in IPF progression (26–28). To test whether TH5487 inhibits relevant lung-resident epithelial and fibroblast cells, A549, BEAS 2B, HFL-1, and murine lung fibroblast cells were included in wound healing assays. TH5487 significantly reduced migration and wound healing compared to TGF-β controls in all tested cell types. In addition, myofibroblast transition is vital to ECM remodeling, leading to lung tissue rigidity and decreased breathing capacity (29). In this regard, HFL-1 myofibroblast transition and proliferation was significantly decreased following TH5487 treatment.

Whilst increased myofibroblast number has been directly linked with IPF progression, fibrotic markers such as collagen, fibronectin, and α-SMA have additionally been linked with ECM deposition and IPF progression (30, 31). In this study, we show that collagen, fibronectin, and α-SMA are reduced by TH5487 treatment in murine lung fibroblast cells stimulated with TGF-β, whilst dexamethasone reduced collagen and α-SMA but importantly did not decrease fibronectin. Fibronectin is a crucial ECM glycoprotein abundantly expressed by fibroblasts, mediating migration and proliferation of myofibroblasts (32). Transition of fibroblasts to myofibroblasts is additionally associated with transcriptional upregulation of profibrotic genes. The effects of TH5487 on transcription of α-smooth muscle actin (α-SMA) and collagen (ColA1) in TGF-β1 stimulated HFL-1 cells were investigated by qPCR. TH5487, 10 μM significantly reduced transcription of Col1A1 and α-SMA while TH5487 at 5 μM reduced mRNA levels of α-SMA in TGF-β1 stimulated cells. Furthermore, immunofluorescence revealed a significant and specific decrease in OGG1 expression following TH5487 treatment, a decrease not seen in TGF-β or dexamethasone controls. Importantly, this confirms the potent and specific activity of TH5487 to decrease OGG1 in fibrosis-inducing cells.

In this study, *in vivo* efficacy of TH5487 was tested using an intratracheal bleomycin model of fibrotic lung damage. Bleomycin-induced fibrosis in C57BL/6 mice is reproducible, only affects the lungs, and is widely used, allowing generation of comparative results (20, 33). Previous studies have shown significant fibrotic-associated weight loss in this model (34), an effect confirmed in this study. Murine weight-loss was significantly higher in groups not receiving TH5487 or dexamethasone, indicative of the protective effects offered by both treatments.

Mechanistic studies using this model, incorporated pro-inflammatory cytokine measurement of various lung environments. BALF, plasma, and lung homogenate displayed significantly decreased levels of a pro-inflammatory and pro-fibrotic cytokines. Specifically, macrophage-recruiting cytokines such as MIP-1α were significantly reduced in the BALF and lung homogenate samples, whilst neutrophil-recruiting MIP-1β (CCL4), KC/CXCL 1, and eotaxin were all reduced following TH5487 treatment.

Interleukin-5 (IL-5) was significantly decreased in the BALF and plasma of TH5487-treated mice. Involved in numerous inflammatory processes in the lungs including IPF (35), IL-5 displays chemoattractant properties for eosinophils, immune cells providing sources of TGF-β and MCP-1 (36, 37). IL-9 was also significantly decreased in both BALF and lung homogenate of TH 5487-treated mice. IL-9 is Th2 cytokine implicated in fibrosis, acting via receptors on macrophages, T/B lymphocytes, and mast cells. High levels of IL-9 have been found in both macrophages and CD4-positive cells in patients with IPF, with IL-9 blockade displaying decreased levels of silica-induced lung fibrosis (38). Additionally, TGF-β1 was shown to be reduced in murine BALF, plasma, and lung homogenate following TH5487 treatment as compared to vehicle/bleomycin treated mice. Elevated TGF-β, a potent inducer of ECM production and key mediator of lung fibrosis, is directly linked to increases in collagen synthesis and deposition (39).

Furthermore, increases in the permeability of the pulmonary vasculature and plasma protein leakage have been linked to the development of pulmonary fibrosis (40). Plasma leakage into the BALF as a marker for lung damage was assessed by the measurement of murine albumin content in the BALF. Murine albumin content was significantly decreased in the BALF following TH5487 treatment as compared to vehicle/bleomycin treated mice indicative of decreased lung damage (41). Taken together, lung injury is accompanied by inflammatory cytokine production, inflammatory cell infiltration, and ensuing fibrosis (42), processes which may be ablated by TH5487 treatment.

To test this, inflammatory cell influx was measured in murine BALF via flow cytometry. Interestingly, fewer neutrophils and significantly fewer inflammatory macrophages were seen in TH5487-treated samples compared to both dexamethasone/bleomycin and vehicle/bleomycin samples. Giemsa-Wright stains of BALF further elucidated the increase in inflammatory macrophage numbers in the bleomycin-treated samples compared to TH5487/bleomycin samples. Inflammatory cell incursion into the lung environment plays a role in IPF progression and severity (16, 43), with limitation of lung inflammation providing a potential source of IPF limitation.

Histological analysis of TH5487 murine lung samples revealed significantly decreased levels of collagen deposition in lung tissues compared to vehicle/bleomycin treated samples. Furthermore, H&E staining of lung samples indicated significant reduction in lung damage and alveolar structural collapse. Further, SEM analysis of TH5487-treated samples revealed decreased thickening of alveolar walls and decreased collagenous deposition compared to vehicle/bleomycin controls. These results provide indicative *in vivo* data of fibrosis reduction following TH5487 treatment. Immunofluorescent staining of fibrotic related proteins revealed decreased levels of both fibronectin and COL1A1 following TH5487 treatment. Additional staining for neutrophil-associated myeloperoxidase (MPO) revealed decreases in neutrophil influx into the lung environment following TH5487 treatment, as supported by similar neutrophil decreases in the BALF flow cytometric analyses. Subsequent co-staining for OGG1/COL1A1 indicates strong signalling of both proteins in similar areas of the murine lung, with both OGG1 and COL1A1 decreased following TH5487 treatment compared to both vehicle/bleomycin and dexamethasone/bleomycin groups.

TH5487 possesses a novel mechanism of action involving upstream gene-targeting of *OGG1* using a small molecule inhibitor. This study further elucidates the downstream effects of this approach, decreasing myofibroblast transition, epithelial and fibroblast migration, inflammatory cell recruitment, and eventual inhibition of fibrotic-related lung remodelling. These data strongly support TH5487 use in clinical trials for IPF treatment.

## Acknowledgement

We would like to acknowledge the assistance received from the IQ Biotechnology Platform (Lund University).

## Authorship contribution statement

**Lloyd Tanner:** Investigation, data curation, writing - review & editing, writing - original draft; **Andrew B. Single:** Investigation, data curation, writing - review & editing; **Ravi K.V. Bhongir:** Investigation, writing - review & editing; **Riya Oomen:** investigation, data curation; **Olov Wallner**: Compound design;**Christina Kalderen**: Conceptualization, funding acquisition; **Thomas Helleday:** Conceptualization, compound design, funding acquisition; **Arne Egesten:** Supervision, conceptualization, funding acquisition, writing - review & editing.

## References

1. Raghu G, Collard HR, Egan JJ, Martinez FJ, Behr J, Brown KK, Colby T V, Cordier J-F, Flaherty KR, Lasky JA, Lynch DA, Ryu JH, Swigris JJ, Wells AU, Ancochea J, Bouros D, Carvalho C, Costabel U, Ebina M, Hansell DM, Johkoh T, Kim DS, King Jr TE, Kondoh Y, Myers J, Müller NL, Nicholson AG, Richeldi L, Selman M, Dudden RF, Griss BS, Protzko SL, Schünemann HJ, Fibrosis AC on IP. 2011. An official ATS/ERS/JRS/ALAT statement: idiopathic pulmonary fibrosis: evidence-based guidelines for diagnosis and management. Am J Respir Crit Care Med 183:788–824.

2. Chang X, Xing L, Wang Y, Yang C-X, He Y-J, Zhou T-J, Gao X-D, Li L, Hao H-P, Jiang H-L. 2020. Monocyte-derived multipotent cell delivered programmed therapeutics to reverse idiopathic pulmonary fibrosis. Sci Adv 6:eaba3167.

3. Raghu G, Flaherty KR, Lederer DJ, Lynch DA, Colby T V, Myers JL, Groshong SD, Larsen BT, Chung JH, Steele MP, Benzaquen S, Calero K, Case AH, Criner GJ, Nathan SD, Rai NS, Ramaswamy M, Hagmeyer L, Davis JR, Gauhar UA, Pankratz DG, Choi Y, Huang J, Walsh PS, Neville H, Lofaro LR, Barth NM, Kennedy GC, Brown KK, Martinez FJ. 2019. Use of a molecular classifier to identify usual interstitial pneumonia in conventional transbronchial lung biopsy samples: a prospective validation study. Lancet Respir Med 7:487–496.

4. Martinez FJ, Collard HR, Pardo A, Raghu G, Richeldi L, Selman M, Swigris JJ, Taniguchi H, Wells AU. 2017. Idiopathic pulmonary fibrosis. Nat Rev Dis Prim 3:17074.

5. Kim HJ, Perlman D, Tomic R. 2015. Natural history of idiopathic pulmonary fibrosis. Respir Med 109:661–670.

6. Landi C, Bergantini L, Cameli P, d’ Alessandro M, Carleo A, Shaba E, Rottoli P, Bini L, Bargagli E. 2020. Idiopathic Pulmonary Fibrosis Serum proteomic analysis before and after nintedanib therapy. Sci Rep 10:9378.

7. Lederer DJ, Martinez FJ. 2018. Idiopathic pulmonary fibrosis. N Engl J Med 378:1811–1823.

8. Mora AL, Rojas M, Pardo A, Selman M. 2017. Emerging therapies for idiopathic pulmonary fibrosis, a progressive age-related disease. Nat Rev Drug Discov 16:755.

9. Guo L, Karoubi G, Duchesneau P, Aoki FG, Shutova M V, Rogers I, Nagy A, Waddell TK. 2018. Interrupted reprogramming of alveolar type II cells induces progenitor-like cells that ameliorate pulmonary fibrosis. NPJ Regen Med 3:1–13.

10. Wang Y, Chen T, Pan Z, Lin Z, Yang L, Zou B, Yao W, Feng D, Huangfu C, Lin C, Wu G, Ling H, Liu G. 2020. 8-Oxoguanine DNA glycosylase modulates the cell transformation process in pulmonary fibrosis by inhibiting Smad2/3 and interacting with Smad7. FASEB J 34:13461–13473.

11. Zhang S, Li J, Li Y, Liu Y, Guo H, Xu X. 2017. Nitric oxide synthase activity correlates with OGG1 in ozone-induced lung injury animal models. Front Physiol 8:249.

12. Pan L, Wang H, Luo J, Zeng J, Pi J, Liu H, Liu C, Ba X, Qu X, Xiang Y, Boldogh I, Qin X. 2019. Epigenetic regulation of TIMP1 expression by 8-oxoguanine DNA glycosylase-1 binding to DNA:RNA hybrid. FASEB J 33:14159–14170.

13. Selman M, King Jr TE, Pardo A. 2001. Idiopathic pulmonary fibrosis: prevailing and evolving hypotheses about its pathogenesis and implications for therapy. Ann Intern Med 134:136–151.

14. Sheppard D. 2015. Epithelial-mesenchymal interactions in fibrosis and repair. Transforming growth factor-β activation by epithelial cells and fibroblasts. Ann Am Thorac Soc 12 Suppl 1:S21–S23.

15. Tang N, Zhao Y, Feng R, Liu Y, Wang S, Wei W, Ding Q, An MS, Wen J, Li L. 2014. Lysophosphatidic acid accelerates lung fibrosis by inducing differentiation of mesenchymal stem cells into myofibroblasts. J Cell Mol Med2013/11/19. 18:156–169.

16. Wynn TA, Ramalingam TR. 2012. Mechanisms of fibrosis: therapeutic translation for fibrotic disease. Nat Med 18:1028–1040.

17. Luo J, Hosoki K, Bacsi A, Radak Z, Hegde ML, Sur S, Hazra TK, Brasier AR, Ba X, Boldogh I. 2014. 8-Oxoguanine DNA glycosylase-1-mediated DNA repair is associated with Rho GTPase activation and α-smooth muscle actin polymerization. Free Radic Biol Med 2014/03/26. 73:430– 438.

18. Visnes T, Cázares-Körner A, Hao W, Wallner O, Masuyer G, Loseva O, Mortusewicz O, Wiita E, Sarno A, Manoilov A, Astorga-Wells J, Jemth A-S, Pan L, Sanjiv K, Karsten S, Gokturk C, Grube M, Homan EJ, Hanna BMF, Paulin CBJ, Pham T, Rasti A, Berglund UW, von Nicolai C, Benitez-Buelga C, Koolmeister T, Ivanic D, Iliev P, Scobie M, Krokan HE, Baranczewski P, Artursson P, Altun M, Jensen AJ, Kalderén C, Ba X, Zubarev RA, Stenmark P, Boldogh I, Helleday T. 2018. Small-molecule inhibitor of OGG1 suppresses proinflammatory gene expression and inflammation. Science (80-) 362:834 LP – 839.

19. Tashiro J, Rubio GA, Limper AH, Williams K, Elliot SJ, Ninou I, Aidinis V, Tzouvelekis A, Glassberg MK. 2017. Exploring animal models that resemble idiopathic pulmonary fibrosis. Front Med 4:118.

20. Tanner L, Single AB. 2019. Animal Models Reflecting Chronic Obstructive Pulmonary Disease and Related Respiratory Disorders: Translating Pre-Clinical Data into Clinical Relevance. J Innate Immun 12:203–225.

21. Lee SB, Kalluri R. 2010. Mechanistic connection between inflammation and fibrosis. Kidney Int Suppl S22–S26.

22. Upagupta C, Shimbori C, Alsilmi R, Kolb M. 2018. Matrix abnormalities in pulmonary fibrosis. Eur Respir Rev 27:180033.

23. Wuyts WA, Agostini C, Antoniou KM, Bouros D, Chambers RC, Cottin V, Egan JJ, Lambrecht BN, Lories R, Parfrey H, Prasse A, Robalo-Cordeiro C, Verbeken E, Verschakelen JA, Wells AU, Verleden GM. 2013. The pathogenesis of pulmonary fibrosis: a moving target. Eur Respir J 41:1207 LP – 1218.

24. Shi L, Dong N, Fang X, Wang X. 2016. Regulatory mechanisms of TGF-β1-induced fibrogenesis of human alveolar epithelial cells. J Cell Mol Med2016/07/15. 20:2183–2193.

25. Hakem R. 2008. DNA-damage repair; the good, the bad, and the ugly. EMBO J 27:589–605.

26. Molina-Molina M, Machahua-Huamani C, Vicens-Zygmunt V, Llatjós R, Escobar I, Sala-Llinas E, Luburich-Hernaiz P, Dorca J, Montes-Worboys A. 2018. Anti-fibrotic effects of pirfenidone and rapamycin in primary IPF fibroblasts and human alveolar epithelial cells. BMC Pulm Med 18:63.

27. Fernandez IE, Eickelberg O. 2012. New cellular and molecular mechanisms of lung injury and fibrosis in idiopathic pulmonary fibrosis. Lancet 380:680–688.

28. King Jr TE, Pardo A, Selman M. 2011. Idiopathic pulmonary fibrosis. Lancet 378:1949–1961.

29. Rangarajan S, Bone NB, Zmijewska AA, Jiang S, Park DW, Bernard K, Locy ML, Ravi S, Deshane J, Mannon RB, Abraham E, Darley-Usmar V, Thannickal VJ, Zmijewski JW. 2018. Metformin reverses established lung fibrosis in a bleomycin model. Nat Med 24:1121–1127.

30. Hu B, Phan SH. 2013. Myofibroblasts. Curr Opin Rheumatol 25:71–77.

31. Phan SH. 2008. Biology of fibroblasts and myofibroblasts. Proc Am Thorac Soc 5:334–337.

32. Beyeler J, Katsaros C, Chiquet M. 2019. Impaired Contracture of 3D Collagen Constructs by Fibronectin-Deficient Murine Fibroblasts.Front Physiol 10:166.

33. Walters DM, Kleeberger SR. 2008. Mouse Models of Bleomycin-Induced Pulmonary Fibrosis. Curr Protoc Pharmacol 40:5.46.1-5.46.17.

34. Egger C, Cannet C, Gérard C, Jarman E, Jarai G, Feige A, Suply T, Micard A, Dunbar A, Tigani B, Beckmann N. 2013. Administration of Bleomycin via the Oropharyngeal Aspiration Route Leads to Sustained Lung Fibrosis in Mice and Rats as Quantified by UTE-MRI and Histology. PLoS One 8:e63432.

35. Boomars KA, Wagenaar SS, Mulder PG, van Velzen-Blad H, Van den Bosch JM. 1995. Relationship between cells obtained by bronchoalveolar lavage and survival in idiopathic pulmonary fibrosis. Thorax 50:1087–1092.

36. Zhang K, Gharaee-Kermani M, Jones ML, Warren JS, Phan SH. 1994. Lung monocyte chemoattractant protein-1 gene expression in bleomycin-induced pulmonary fibrosis. J Immunol 153:4733 LP – 4741.

37. Zhang K, Flanders KC, Phan SH. 1995. Cellular localization of transforming growth factor-beta expression in bleomycin-induced pulmonary fibrosis. Am J Pathol 147:352–361.

38. Sugimoto N, Suzukawa M, Nagase H, Koizumi Y, Ro S, Kobayashi K, Yoshihara H, Kojima Y, Kamiyama-Hara A, Hebisawa A, Ohta K. 2018. IL-9 Blockade Suppresses Silica-induced Lung Inflammation and Fibrosis in Mice. Am J Respir Cell Mol Biol 60:232–243.

39. Yue X, Shan B, Lasky JA. 2010. TGF-β: Titan of Lung Fibrogenesis. Curr Enzym Inhib 6:10.2174/10067.

40. Shea BS, Probst CK, Brazee PL, Rotile NJ, Blasi F, Weinreb PH, Black KE, Sosnovik DE, Van Cott EM, Violette SM, Caravan P, Tager AM. 2017. Uncoupling of the profibrotic and hemostatic effects of thrombin in lung fibrosis. JCI insight 2:e86608.

41. Knipe RS, Probst CK, Lagares D, Franklin A, Spinney JJ, Brazee PL, Grasberger P, Zhang L, Black KE, Sakai N, Shea BS, Liao JK, Medoff BD, Tager AM. 2018. The Rho Kinase Isoforms ROCK1 and ROCK2 Each Contribute to the Development of Experimental Pulmonary Fibrosis. Am J Respir Cell Mol Biol 58:471–481.

42. Phan SH, Kunkel SL. 1992. Lung Cytokine Production in Bleomycin-Induced Pulmonary Fibrosis. Exp Lung Res 18:29–43.

43. Butler MW, Keane MP. 2019. The Role of Immunity and Inflammation in IPF Pathogenesis BT - Idiopathic Pulmonary Fibrosis: A Comprehensive Clinical Guide, p. 97–131. In Meyer, KC, Nathan, SD (eds.). Springer International Publishing, Cham.

