## Supplementary Figures 1-5 for "Small-molecule-mediated OGG1 inhibition attenuates pulmonary inflammation and lung fibrosis"

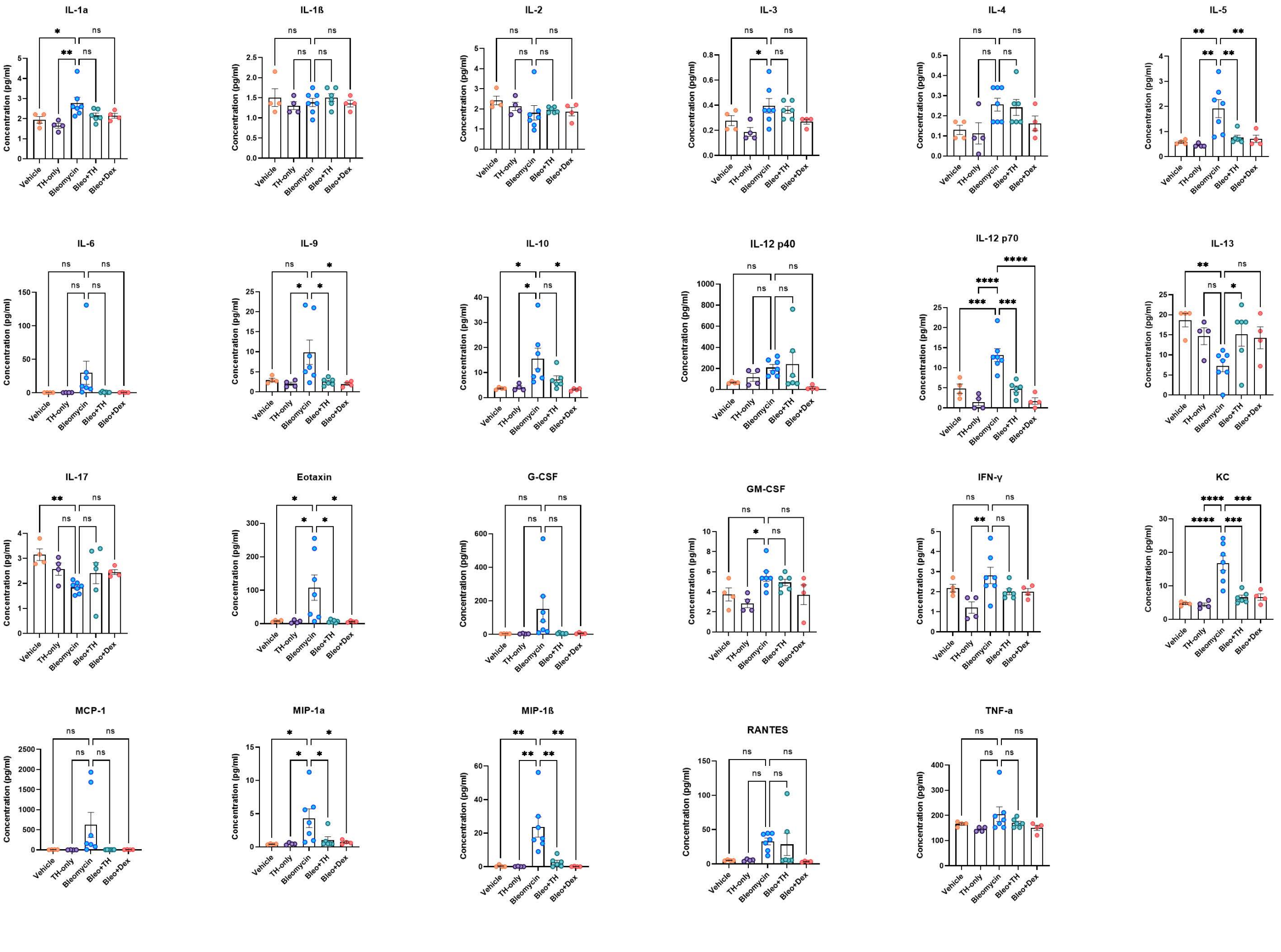


**Supplementary Figure 1: Murine BALF cytokine levels.** Cytokine values were compared to the vehicle/bleomycin group using a one-way ANOVA (**P<0.05; **P<0.01; ***P<0.005; *****P<0.001*).

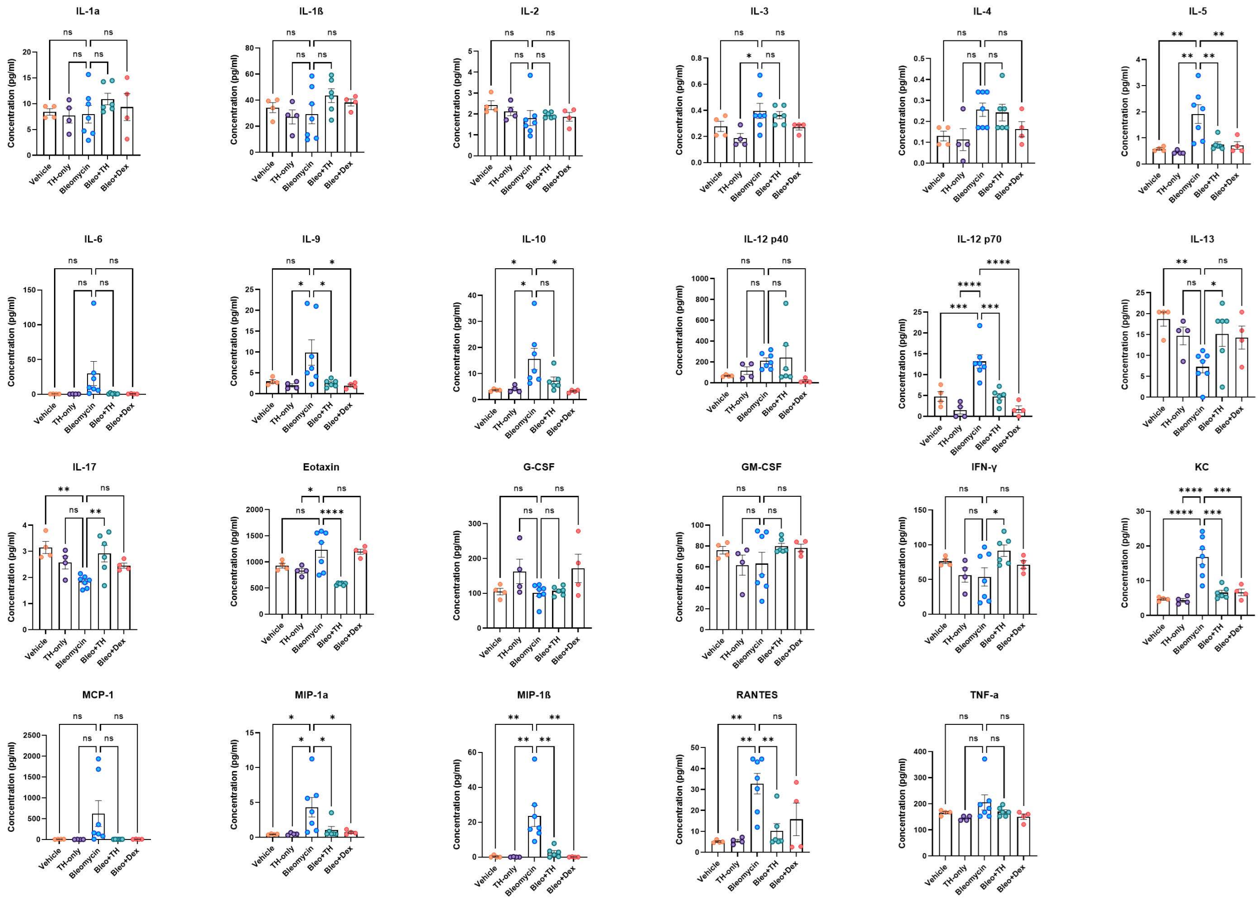


**Supplementary Figure 2: Murine plasma cytokine levels.** Cytokine values were compared to the vehicle/bleomycin group using a one-way ANOVA (**P<0.05; **P<0.01; ***P<0.005; *****P<0.001*).

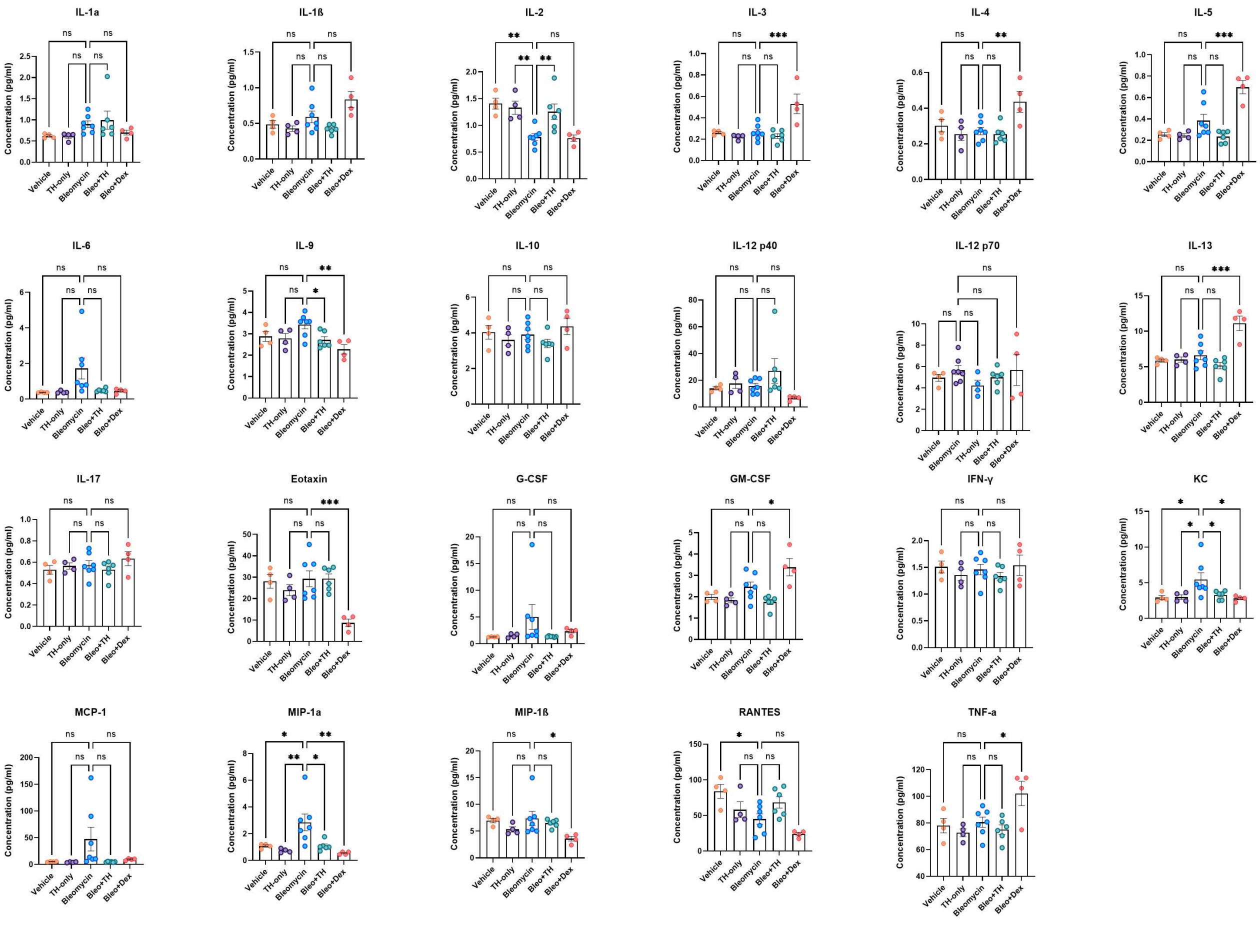


**Supplementary Figure 3: Murine lung homogenate cytokine levels.** Cytokine values were compared to the vehicle/bleomycin group using a one-way ANOVA (**P<0.05; **P<0.01; ***P<0.005; *****P<0.001*).

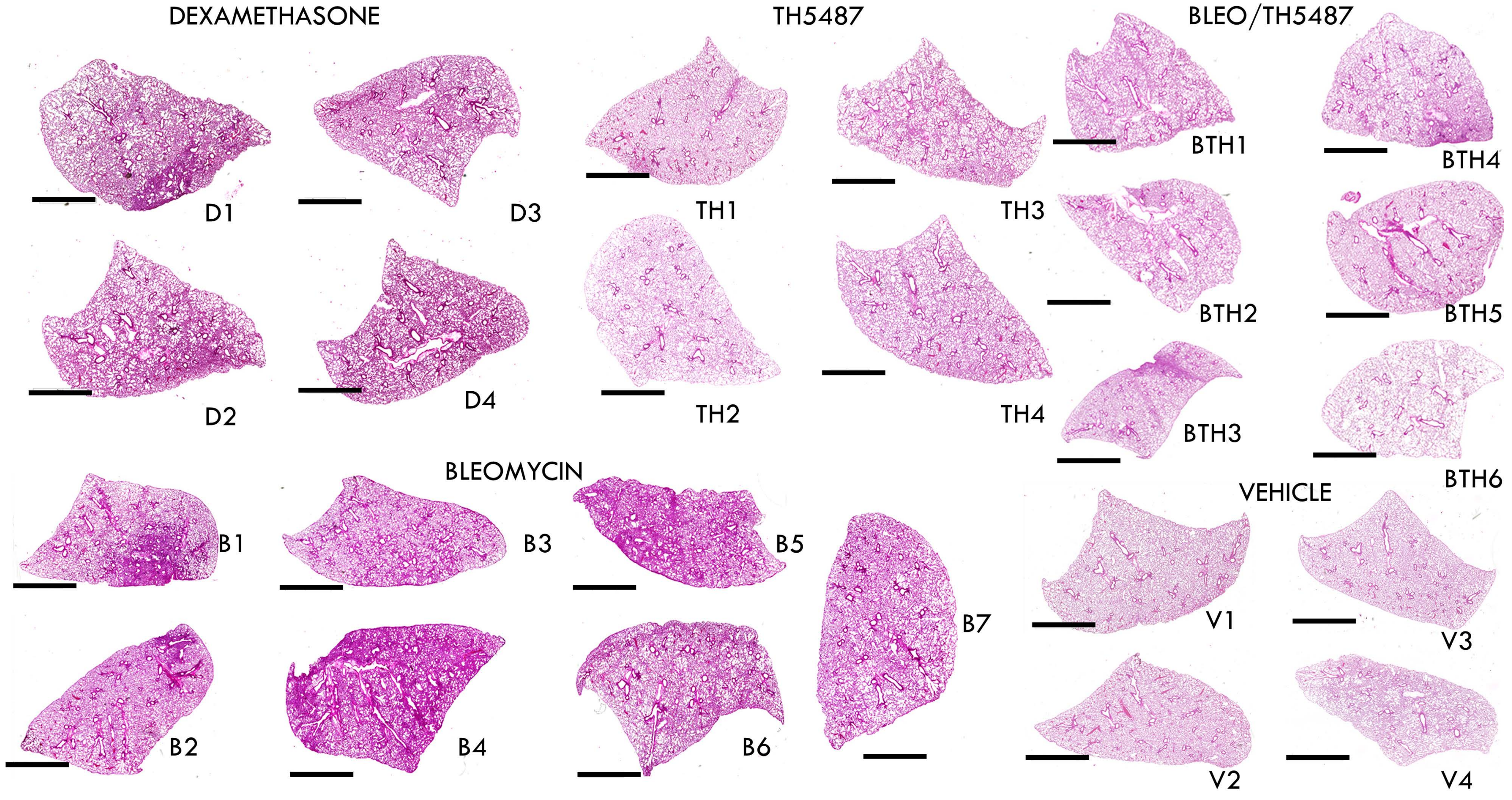


**Supplementary Figure 4: Whole lung scans of murine lungs following H&E staining (scale bar=2 mm).**

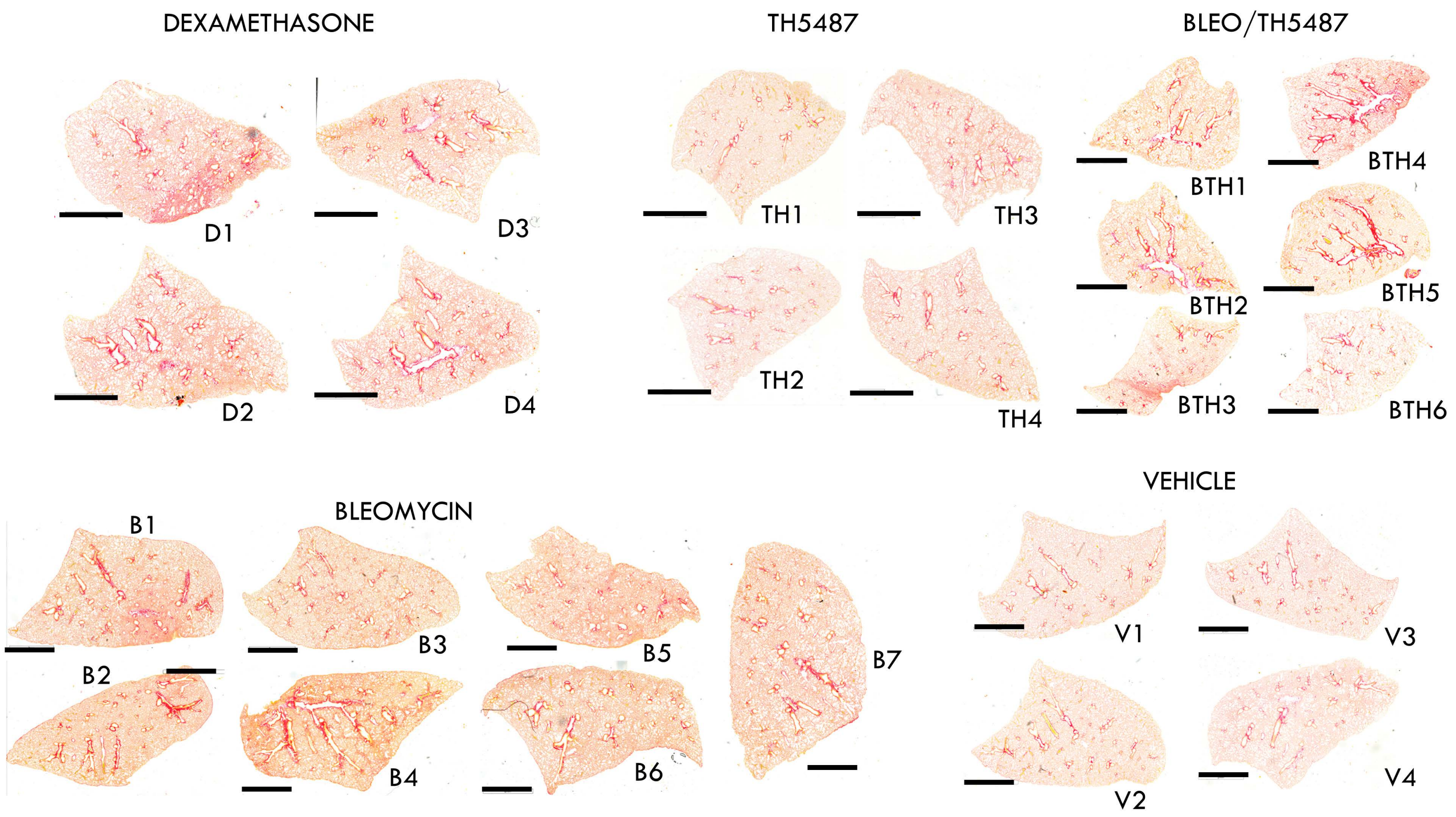


**Supplementary Figure 5: Whole lung scans of murine lungs following picrosirius red staining (scale bar=2 mm).**
